# Ingestible Osmotic Pill for *In-vivo* Sampling of Gut Microbiome

**DOI:** 10.1101/690982

**Authors:** Hojatollah Rezaei Nejad, Bruno C. M. Oliveira, Aydin Sadeqi, Amin Dehkharghani, Ivanela Kondova, Jan A.M. Langermans, Jeffrey S. Guasto, Saul Tzipori, Giovanni Widmer, Sameer R. Sonkusale

## Abstract

Technologies capable of non-invasively sampling different locations in the gut upstream of the colon will enable new insights into the role of organ-specific microbiota in human health. We present an ingestible, biocompatible, battery-less, 3D-printed micro-engineered pill with integrated osmotic sampler and microfluidic channels for *in vivo* sampling of the gut lumen and its microbiome upstream of the colon. The pill’s sampling performance was characterized using realistic *in vitro* models and validated *in vivo* in pigs and primates. Our results show that the bacterial populations recovered from the pills’ microfluidic channels closely resemble the bacterial population demographics of the microenvironment to which the pill was exposed. We believe such lab-on-a-pill devices will revolutionize our understanding of the spatial diversity of the gut microbiome and its response to medical conditions and treatments.

## Introduction

The gut microbiota represents trillions of bacteria belonging to around one thousand species.^[1]^ Among the various microbial communities associated within the human body, the gut microbiome is noted for its diversity, its elevated concentration^[2,3]^ and its many beneficial functions. Dysbiosis (microbial imbalance) has been associated with conditions such as inflammation, recurrent infections and increased susceptibility to enteric pathogen.^[4–8]^ Most studies infer the condition of the gut microbiome from the analysis of fecal DNA and fecal metabolites.^[1]^ Because the gut environment changes as the gut content moves down the gastrointestinal (GI) tract, analyses of feces are inadequate to identify abnormal conditions upstream of the distal colon. Whereas the analysis of complex bacterial populations has benefitted from new DNA sequencing techniques, our capacity to precisely and non-invasively sample different organs has not improved. Consequently, medical research is often based on easily accessible samples, like feces, in spite of the inherent limitations of the conclusions, which can be drawn from the analysis of such samples. For instance, samples important for understanding the interaction between enteric pathogens and the host remain out of reach, unless invasive sampling techniques are used. To gain new insights into the many beneficial functions of the gut microbiota, it is essential to sample *in vivo* different locations in the gut, particularly organs located upstream of the colon.

Here, we describe the development and testing of an innovative non-invasive technology to sample the intestinal lumen *in vivo.* We created an ingestible, biocompatible, 3D-printed microengineered pill with integrated osmotic sampler that requires no battery for its operation. Stereo-lithography (SLA) based 3D printing was used to fabricate the miniaturized ingestible device with sophisticated microfluidic functions for spatial sampling of the gut lumen. The pill is covered with a pH sensitive enteric coating to delay sampling until the pill has entered the small intestine, where the coating dissolves in the higher pH environment. A magnetic holding mechanism was designed to enable the pill to sample more time from a targeted region of the gut. Natural peristaltic motion endows mobility to the pill through the GI tract without any active parts. The sampling function of the pill has been extensively validated *in vitro* and *in vivo* in pigs and primates. The magnetic hold for spatial targeting has been validated *in vitro.*

## Results and discussion

### Osmotic pill sampler design

The overall design of the pill is shown in Figure 1a. The osmotic pill sampler consists of three main parts – a top sampling head, a semi-permeable membrane in the middle and a bottom salt chamber. The head comprises four inlets, a stilling basin connected to four helical channels, all of them leading to one small chamber. The salt chamber consists of a cavity that contains dry calcium chloride salt powder, a second cavity to hold a small neodymium magnet, two tube-like reservoirs to hold fluorescent dye, and a horn-shaped exit nozzle at the bottom. The pill works on the principle of osmosis where pressure differential is created across the semipermeable membrane, which creates a passive pumping action (see Figure 1a). This mechanism facilitates the flow of water across the membrane from the helical channels towards the salt chamber. The porosity of the membrane blocks the flow of larger particles (e.g., microorganisms), leaving them trapped in the helical channels (Figure 1a).

**Figure 1.**
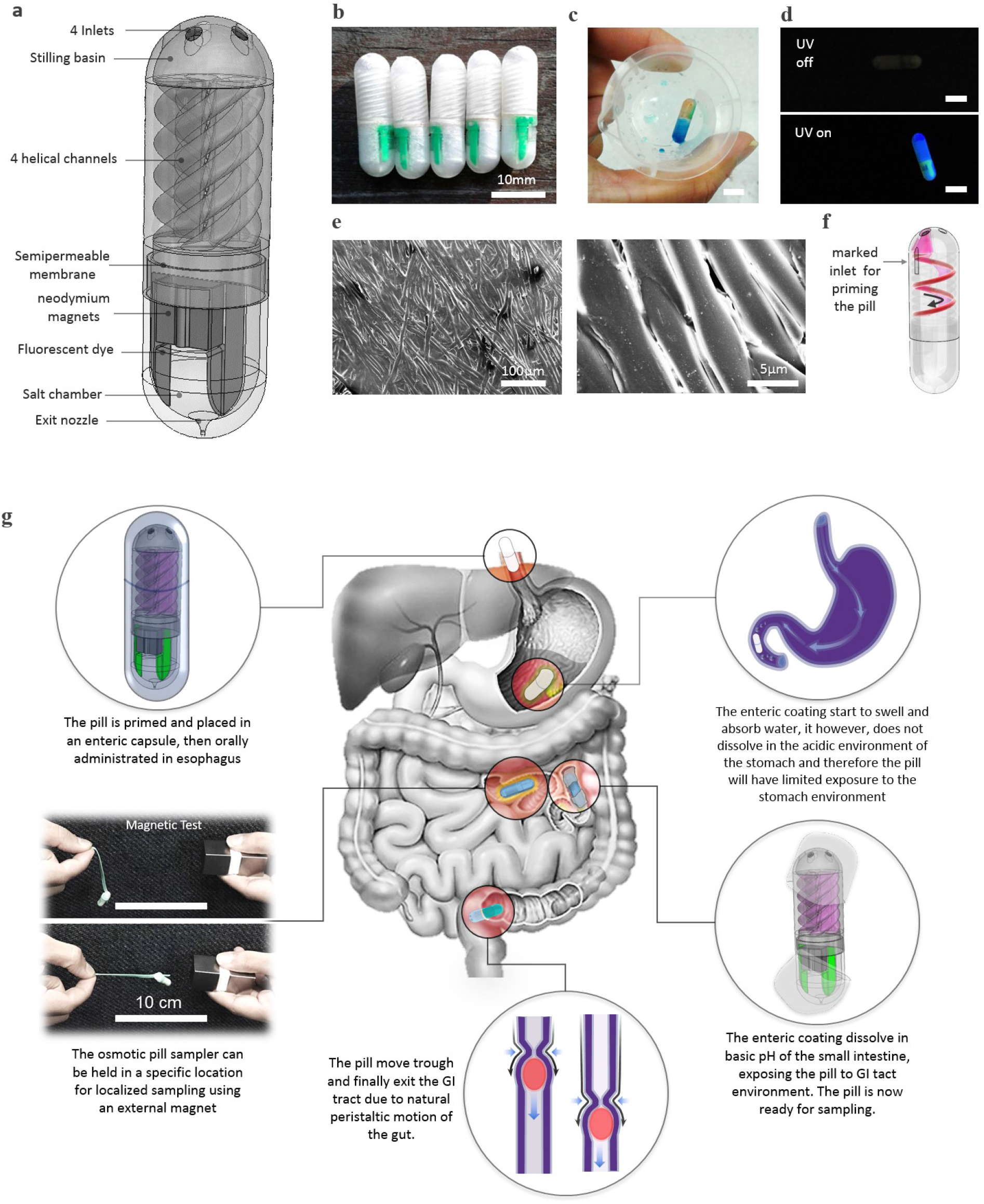
Osmotic pill sampler design and working principle. a) 3D schematic of the overall design of the pill, b) fabricated osmotic pills samplers, c) proof of concept, the pill samples a water solution containing blue food dye (the sample did not penetrate into the salt chamber), d) the pill surface and the fluorescent dye in the pill are detectable under UV light, e) Scanning Electron Microscope (SEM) images of the osmotic membrane used in the pill, f) marked inlet at the top chamber is designed for initiating the pill (priming), g) overall working principle of the osmotic pill sampler in an enteric capsule.

The fabrication of this pill is explained in detail in the Materials and Methods section. Briefly, the pill sampling head and the bottom salt chamber are 3D printed using a stereo-lithography 3D printer. A semipermeable membrane is used to construct the osmotic pump. The membrane itself is made of woven 5 μm thick cellulose acetate fibers commonly used in reverse osmosis based water filters (Figure 1e,f). A small neodymium magnet is placed and sealed inside the salt chamber. To facilitate locating the pill after it is excreted in feces, two lines of fluorescent green dye are painted and sealed in the salt chamber. The salt chamber is filled with calcium chloride salt. The top and bottom chambers are separated by the semipermeable membrane assembled and attached together using UV curable adhesive. The fabricated osmotic pill sampler is shown in Figure 1b.

### Pill operation concept

#### Priming the pill

The pill is primed by injecting approximately 200 μL water into the salt chamber through the exit nozzle using a syringe with a 36-gauge needle. Similarly, the helical channels are filled with water from a marked inlet, which is connected to one of the helical channels (see Figure 1a, and supplementary video 1). By aspirating water through the marked inlet, the water flows first through the corresponding channel and then fills the chamber at the bottom of the sampling head, and finally fills all the other helical channels and the stilling chamber on the top of the pill in this sequence. The head can hold approximately 120 μL of sample.

#### Sampling strategy

After priming, the pill is ready for oral administration. To avoid sampling the stomach lumen and limit the sampling to the small and large intestine, we placed the pill in a commercially available size 0 enteric coating capsule. The enteric coating resists the acidic environment of the stomach and only dissolves in the neutral/basic environment of the intestine. The dissolution profile for the enteric coating can be controlled by an appropriate choice of polymer(s). Testing of different enteric coatings was not the focus of this study. The osmotic pill sampler moves down the GI tract primarily due to natural peristalsis. The pill, however, can be held at a specific location inside the GI tract using an external magnet. This is made possible by a small neodymium magnet embedded in the pill (Figure 1a). Immobilizing the pill enables preferential sampling of specific regions of the gut, so that more of the collected sample originates from this region. Without this magnet, the pill would sample more or less uniformly along the entire length of the GI tract. We have added two fluorescent marks on the pill to facilitate detection following excretion (Figure 1d). The details of our sampling strategy are shown in Figure 1g.

### Physical characterization of the pill

Osmosis is a net movement of a solvent through a semi-permeable membrane towards a region of high solute concentration. Once the pill is primed, the process of osmosis causes solvent (water) to flow across the membrane from the helical channels into the salt chamber (Figure 1f). The flow rate depends on the properties of this membrane (e.g. thickness, porosity, area) and the salt gradient across it. The fluid sampler consists of such an osmotic pump that continuously pulls the fluid from the gut into the narrow microfluidic collection channels (<1 mm diameter) at a flow rate of 2 ~ 20 μL/hr. We characterized the flow created by osmosis across the semipermeable membrane, which drives the sampling function of the pill (see Supplementary section 1a). The flow rate through the membrane assembled in the pill was also measured over time (Supplementary Figure 1b). Initially, the flow rate was ~2.5 μL/hr and remained approximately constant for almost 48 hr. The flow rate fell to around ~0.5 μL/hr after four days.

The design of the exit nozzle in the salt chamber is important because it facilitates the discharge of water being accumulated in the salt chamber as a result of the osmotic action. Numerical simulations were carried out to study flow through the discharge hole for two different hole diameters, 100 μm and 50 μm (see supplementary Figure 2). The maximum fluid velocity for the 50 μm hole is 0.6 mm/s, which is 4.28 times higher than for hole of 100 μm size. A higher continuous flow rate at the discharge point has two advantages: first, it reduces the diffusion from the environment into the salt chamber, essentially acting as a one-way valve; second, a high flow rate reduces the chance of blockage of the discharge hole by solid debris present in the gut.

### *In vitro* sampling

The gut microbiome comprises prokaryotic and eukaryotic cells that may have no active motility or may be able to swim propelled by flagella or other mechanisms. We validated whether the osmotic pump can sample microorganisms regardless of their motion by testing the pill *in vitro* with non-motile solid particulates as well as with highly motile bacteria. The first environment included polystyrene micro-particles of 10μm diameter in a highly viscous liquid (160 mPa.s) to mimic the gut environment. These particles exhibit negligible diffusion, especially in the highly viscous solution of 50% w/v polyethylene glycol hydrogel. Since these particles are neutrally buoyant in this solution, the effect of gravity on their motion is insignificant. The second environment included highly motile bacteria, wild-type *Bacillus subtilis,* in an aqueous solution. These bacteria perform a run-and-tumble motion,^[9]^ meaning they swim in a certain direction for a short time (~1 s) and then change their swimming direction at random. This results in a highly diffusive transport of the bacteria over long times with effective diffusion coefficients comparable to small gas molecules in water.

### *In vitro* sampling of non-motile micro-particles

We tested the pill in highly viscous environments similar to the GI tract using solutions of 50% (w/v) of polyethylene glycol (PEG), with a molecular mass of 10,000, in two different aqueous buffers of pH 4 and pH 8. Different pH conditions help model the fact that pH varies along the GI tract. The measured viscosity of the solution was 160 ±5 mPa.s at 25°C (USS-DVT4 Rotary Viscometer Viscosity Meter, U.S. Solid Inc., Cleveland, OH, USA). Next, polystyrene particles 10 μm in diameter were added to these solutions to mimic partially digested food particles in the GI tract. These micro-particles are large enough so that the Brownian motion can be ignored. Therefore, any micro-particles captured by the pill would have entered the helical channels mainly be due to the osmotic flow. Initially, pills were primed with deionized water without PEG. Then, a group of three primed pills were placed in each of the pH 4 and pH 8 solutions described earlier. At different time points, a pill was taken out of the solution, and the sample was extracted from the pill using a pipette. On average 120 ± 6 μL sample was recovered from each pill. The extracted sample was weighed and imaged using an optical microscope and finally dried on a hotplate at 40°C and weighed again. The samples were weighed before and after drying to evaluate the amount of hydrogel aspirated by the pill. The number of micro-particles acquired by the pill at two different time steps was also quantified (see Materials and Methods section for details). To exclude the effect of background fluid motion on the sampling capability of the pill, the tests were performed in static condition with no agitation. To justify exclusion of the uptake in the pill by diffusion, we calculate the diffusion coefficient, *D*, for micro-particles in the viscous medium through the Stokes-Einstein relation,

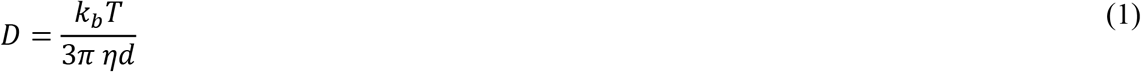

where *k_b_* is Boltzmann’s constant, *T* is the absolute temperature, *d* is the diameter of the particle, and *η* is the dynamic viscosity of the medium. The diffusion coefficient of a 10 μm diameter spherical particle at 37°C in a fluid with viscosity of 160 mPa.s is *D* = 2.8 × 10^-4^ μm^2^/s. To diffuse one particle diameter under such conditions would require a time *t* ≈ *d*^2^/2*D* = 2.1 days. Thus, on time scales relevant to our experiments, diffusion is ineffective in generating any appreciable uptake of particles into the pill.

Figure 2 shows the number of particles recovered from the helical channels for both acidic (pH 4) and basic (pH 8) environments. The number of particles increases over time (Figure 2a), which validates that the osmotic pill can indeed sample non-motile micro-particles in a highly viscous environment. We also evaluated the amount of PEG aspirated into the pill’s helical channels over time. Initially, the channels contain only pure water used for priming. Over time, the PEG from the surrounding environment is sampled by the pill along with micro-particles, causing the concentration of the PEG in the pill to increase. Figure 2b shows a quantitative comparison of the hydrogel concentration in the pill over time in acidic (pH 4) and basic (pH 8) conditions. The hydrogel concentration increases almost linearly with time. However, in basic conditions, a higher sampling rate was noticed, as was a higher hydrogel concentration inside the pill, exceeding the concentration in the surrounding environment. We believe this is because the pH protonates and deprotonates the functional groups of the membrane itself and of the molecules in solution. This effect will change the effective charge on the membrane and alter the size of the pores, which will impact the membrane’s nanofiltration properties and affect the flux of water.^[10,11]^ The zeta potential (ζ) of a cellulose acetate membrane is negative for pH 3 or higher and becomes more negative as the pH increases.^[12]^ Therefore, the membrane is more negatively charged at pH 8 (ζ ≈ −35 mV) compared to pH 4 (ζ ≈ −10 mV), which can result in bigger pore sizes and higher flux at pH 8. We repeated these experiments for a total of 12 pills; six under acidic (pH 4) conditions and the rest at pH 8. The same 50% PEG (w/v) solution and black 10 μm polystyrene beads of were used as discussed earlier. The results shown in Figure 2 validate the sampling capability of the pill in both acidic and basic conditions. The number of particles is larger under basic conditions as compared to the acidic conditions for the reasons explained earlier, and is approximately equal to the control after 24 h of operation. Here, the control is the liquid containing the particles in which pills are immersed.

**Figure 2.**
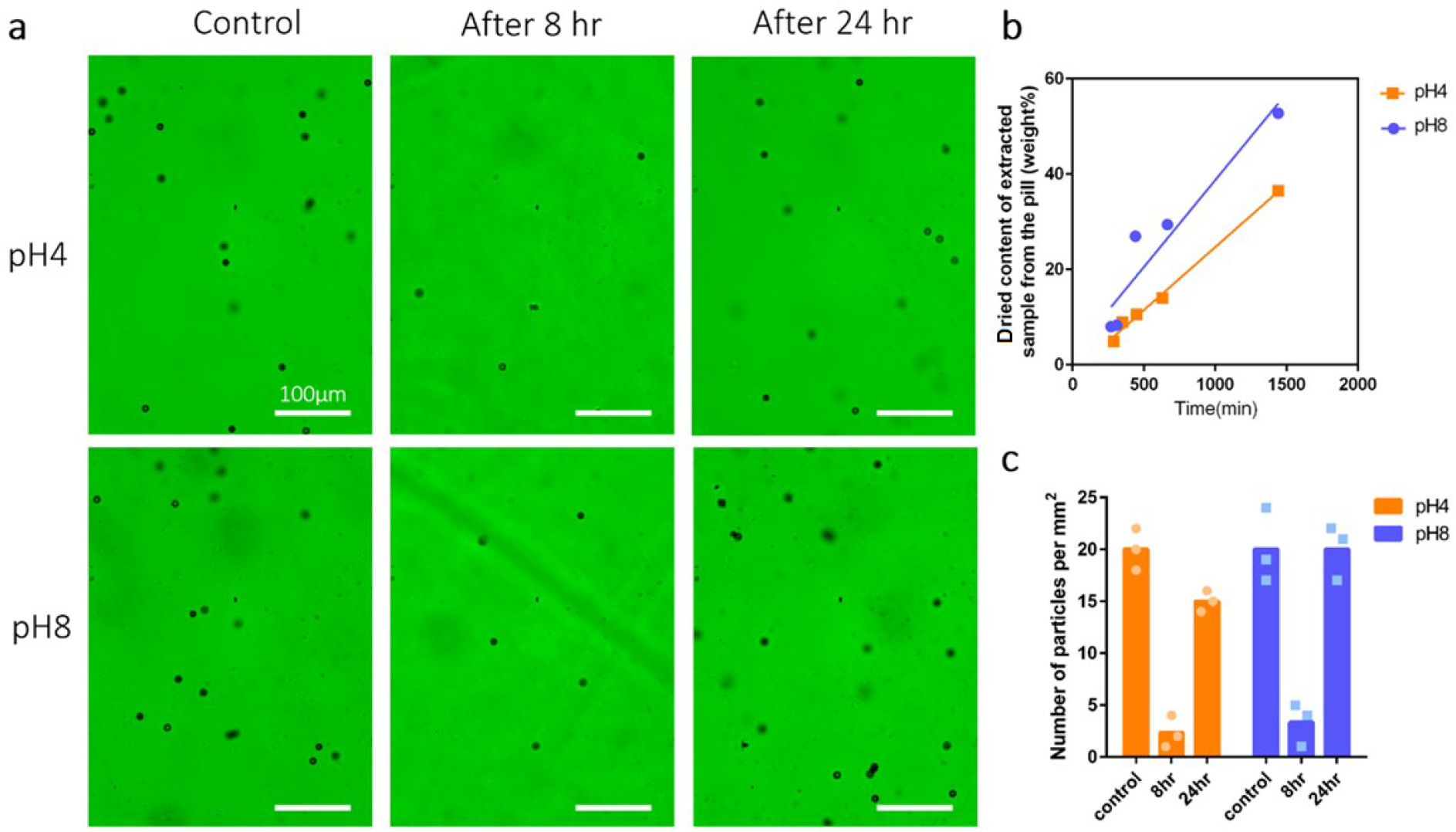
Osmotic sampling by the pill of solid particles in acidic and basic high-viscosity environment (50% polyethylene glycol hydrogel). a) micro-particles (polystyrene beads) captured by the pill after 8 hr and 24 hr in solutions of different pH. Control shows particle suspension in which pills were immersed. b) weight percentage of the dried sample extracted from the pill at different time points. c) number of particles captured by the pill in two different pH conditions and compared to control (control is considered the environment outside of the pill).

### *In vitro* sampling of motile bacteria

In the second *in vitro* test, we aimed to evaluate the sampling performance of the osmotic pills with motile bacteria. Wild-type *B. subtilis* was chosen as the test bacterium, which is ubiquitous in the GI tract of ruminants and humans. We used a fluorescent strain to facilitate imaging. *B. subtilis* is a multiflagellated bacterium that swims with an average speed of 52 μm. s^-1^ at room temperature (25°C) when the flagella rotate in synchrony to form a bundle.^[13]^ Fluctuations in flagellar rotation can cause the flagella to unbundle and generate a random reorientation of the cells. Subsequently, the bundle reforms resulting in the canonical run-and-tumble motion of the bacteria, which is characterized by an effective diffusion coefficient,^[14]^

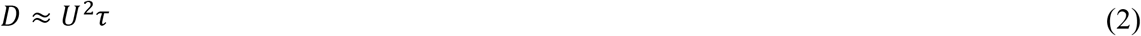

where, *U* is the swimming speed of the cells and *τ* is the persistence time of the cell orientation, which is approximately ~1 s for *B. subtilis*.^[15]^ Therefore, the effective diffusion coefficient is 2700 μm^2^/s, which is 10^7^ larger than the polystyrene beads used in the previous experiment.

Figure 3 shows the motile bacteria sampled by the pill. Nine osmotic pills were placed in separate test tubes containing the bacterial suspension, and three pills removed at two time points. Their content was examined under a fluorescent microscope and compared with a sample of bacterial suspension (see Materials and Methods for details). The concentration of bacteria in the pill increased over time, and after 1.5 hr it was almost equal to the concentration in the environment. This outcome was in fact expected as these bacteria are very motile and their diffusion factor is orders of magnitude higher than passive particles of the same size. Significantly, after 3 hr, the bacteria in the pill were almost three times more concentrated than in the surrounding environment, likely the effect of continuous osmotic sampling. The osmotic membrane is water permeable, which in essence is filtering the bacteria and accumulating them in the collection chamber. Moreover, the helical geometry of the channels reduces the probability that motile bacteria can leave the collection chambers. The results also take into account bacterial replication through the control (the surrounding medium outside of the pill), and we see no increase in bacterial concentration in the suspension and definitely cannot justify a 3-fold increase in the concentration after 3 hr. The *in vitro* test was undertaken for 24 hr. As expected, the bacteria captured by the pill remained motile.

**Figure 3.**
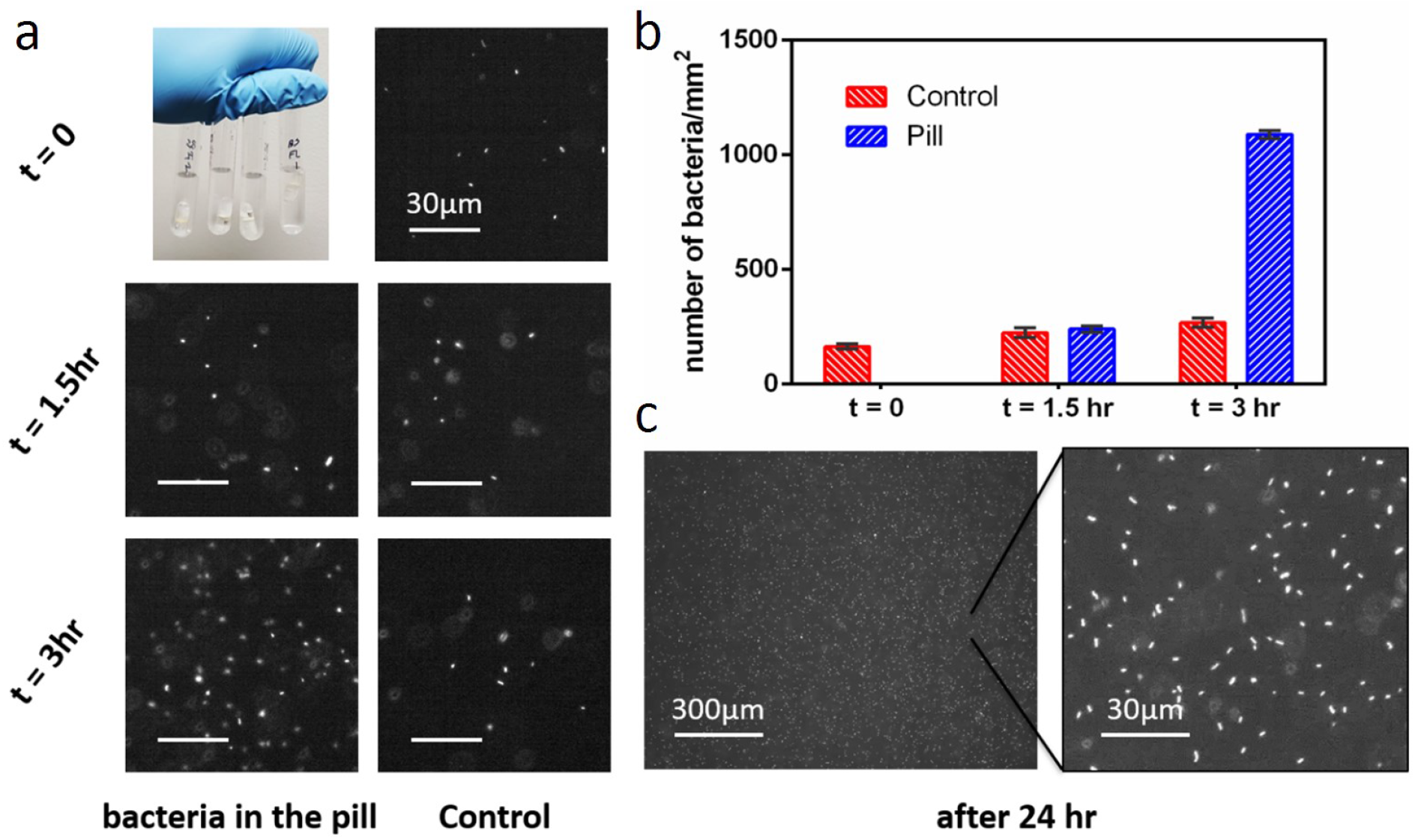
Osmotic sampling by the pill of motile bacteria in aqueous suspension (medium). Bacillus subtilis was used to model live microorganisms. a) snapshot of bacteria concentration in the pill and surrounding environment outside of the pill (control) at different time points. b) number of bacteria sampled by the pill and outside of the pill (control). The error bars shows the standard deviation of 3 replicate counts of the same sample. c) snapshot of bacteria in the pill after 24 hr.

### Pill mobility in the gut (ex-vivo study)

To study how the pill moves inside the GI tract, we conducted ex-vivo experiments using intestines freshly dissected from pigs (Supplementary Figure 3). At a flow rate of 4 mL/min and 8 mL/min the pills moved with an average speed of ~10 cm/min and ~13 cm/min, respectively. The pill, however, moves with much higher velocity (~22 cm/min) when the flow rate is increased to 10 mL/min, demonstrating nonlinear behavior attributed to the viscoelastic nature of the intestine which expands under increased peristaltic pressure. The natural peristaltic flow rate of fluid through the proximal small intestine varies widely from an average of 2.5 mL/min in fasting subjects to as high as 20 mL/min after meals.^[16–19]^ The *ex vivo* observation indicates that the pill readily moves inside the GI tract under realistic flow conditions.

### *In vitro* and *in vivo* studies

To further investigate the performance the pill, three experiments were performed *in vitro* (see Materials and Methods). In the first experiment, pills were immersed sequentially in suspensions of *Escherichia. coli, B. subtilis* and *Lactobacillus rhamnosus* or only in *E. coli* and *L. rhamnosus.* DNA extracted from the material collected by the pills over a 12-hr immersion period was analyzed by 16S ribosomal RNA (rRNA) amplicon sequencing to quantify the relative abundance of each species and monitor the activity of the osmotic pump over time. Pill 1 was immersed for 4 hr each in cultures of *E. coli*, *B. subtilis* and *L. rhamnosus*. Pills 2 and 3 are exact replicates and were immersed 8 hr in *E. coli* followed by a 4-hr immersion in *L. rhamnosus* culture. Since the number of rRNA operons varies by species and strain, the number of 16S sequences assigned to each species was divided by the number of operons in the genome of each strain used in the experiment. The normalized relative abundance values shown in Table 1 are thus proportional to the number of bacteria in the sample. These data indicate that bacteria sampled in the latter part of each experiment tend to be over-represented. This conclusion is based on the observation that *L. rhamnosus* sequences (order Lactobacillales) are more abundant than expected based on the duration of immersion in *L. rhamnosus* culture. The three rightmost columns of Table 1 show that the vast majority (>90%) of 16S sequences obtained from pure cultures of each bacterium are correctly classified.

**Table 1.**
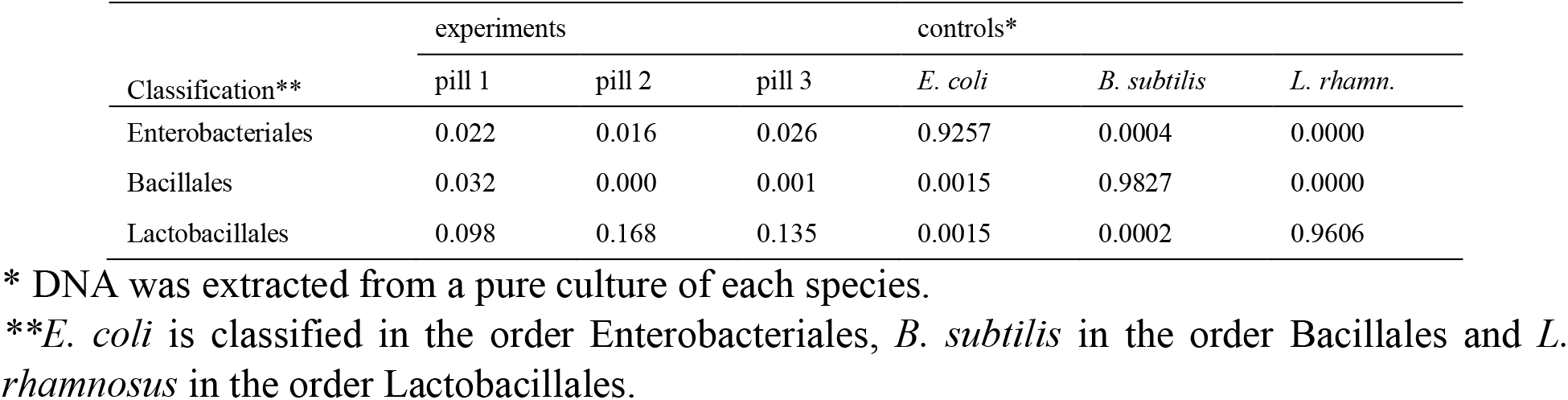
Normalized abundance of three bacterial species in samples recovered from collection channels in experiment 1

| Classification** | experiments |  |  | controls* |  |  |
| --- | --- | --- | --- | --- | --- | --- |
|  | pill 1 | pill 2 | pill 3 | *E. coli* | *B. subtilis* | *L. rhamn.* |
| Enterobacteriales | 0.022 | 0.016 | 0.026 | 0.9257 | 0.0004 | 0.0000 |
| Bacillales | 0.032 | 0.000 | 0.001 | 0.0015 | 0.9827 | 0.0000 |
| Lactobacillales | 0.098 | 0.168 | 0.135 | 0.0015 | 0.0002 | 0.9606 |

\* DNA was extracted from a pure culture of each species.
\*\**E. coli* is classified in the order Enterobacteriales, *B. subtilis* in the order Bacillales and *L. rhamnosus* in the order Lactobacillales.

A second *in vitro* experiment was conducted to confirm that uptake of sample by the pill requires the action of the osmotic pump. Standard pills with CaCl_2_ in the salt chamber as shown in Figure 1a and control pills lacking salt were immersed in parallel for 12 hr in fecal slurry prepared by homogenizing a volume of approximately 5 ml of cat or mouse feces into 500 ml of distilled water. Bacterial DNA extracted from each pill’s collection channels was quantified using quantitative real-time PCR as described in Materials and Methods. Confirming that the action of the osmotic pump is necessary for sampling, this analysis showed that after a 12-hr immersion in fecal slurry the concentration of bacterial DNA in the no-salt pills was 2.4 to 59-fold more dilute than in the regular pills.

The ability of the pill to sample gut luminal content was tested in a weaned pig and in 4 adult rhesus macaques *(Macaca mulata).* A Principal Coordinates analysis (PCoA) combining the 16S sequence data from pig and *in vitro* experiments is shown in Figure 4. As expected, a clear clustering of the microbiota by sample and by organ was observed; pig stomach/jejunum, pig colon/feces, cat and mouse fecal slurry generated well-defined clusters. In the *in vivo* experiments, the microbiota profile of samples recovered from the pills’ collection channels closely resembled the profile of samples recovered from the surrounding intestinal lumen or feces (Supplementary Table 1). Based on this observation, we tested in *in vitro* experiment 3 to what extent the microbiota profile was impacted by material adhering to the surface of the pill, as opposed to material collected in the helical channels. To unequivocally demonstrate that the pills’ osmotic pump actively aspirated material from the surrounding environment into the channel, pills from the *in vitro* experiment were briefly transferred from cat (or mouse) fecal slurry to mouse (or cat) slurry. Figure 4 shows that the samples recovered from the collection channels, designated with a circle in the figure and labeled “in” in the key, indeed originated from the fecal slurry in which the pills were immersed for 24 hr. (triangles; “out”), and that the slurry from the opposite species did not contaminate the sample to a detectable extent. This is apparent from the tight clustering of the red (mouse) triangle with the circle of the same color, and the green (cat) triangle with the green circles. The 16S sequence data show that the pills operate as intended and are capable of actively collecting sample from the surrounding matrix.

**Figure 4.**
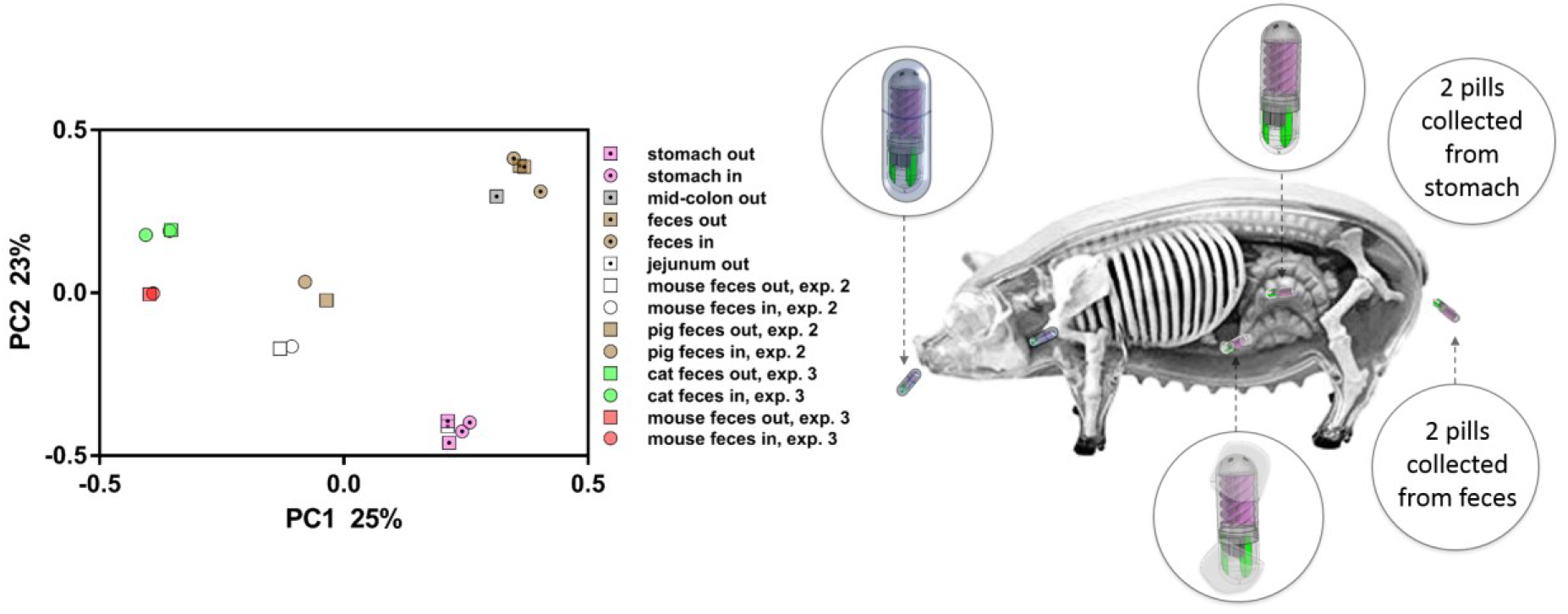
Principal Coordinate Analysis of bacterial populations sampled with pills. Results from two experiments in pigs and two experiment in vitro (experiments 2 and 3) are represented with dotted and empty symbols, respectively. Symbol shape indicates whether sample originated from the pill’s collection channel (○, “in”) or was collected from the matrix from which the pill was recovered (□, “out”). Color indicates organ for in vivo experiments and species origin of fecal sample for in vitro experiments. Two different mouse feces were used in experiment 2 and 3, explaining the distance between white and red datapoints.

Primed pills fitted with enteric coating were also administered individually to 8 adult rhesus macaques. With this experiment we wished to test the functionality of the pill in an animal model with a GI anatomy closely resembling that of humans. Of the 8 pills gavaged to macaques, 3 were successfully recovered from the feces within 2 days of administration. For 5 animals, no pill was found in the feces. None of the missing pills were localized upon post-mortem examination of the GI tract or CT scanning, indicating that the pills were excreted but were missed when the feces were searched.

PCoA of 16S sequences from pills excreted by the (primates), as well as from matching fecal samples are shown in Figure 5. The position on the plot of the samples recovered from the pills’ collection channels is clearly distinct from the feces collected directly from the ground or from feces adhering to the outside of the pills. These results indicate that the pills channels contained sample collected from more proximal organs, consistent with the pills’ ability to sample the entire GI tract.

**Figure 5.**
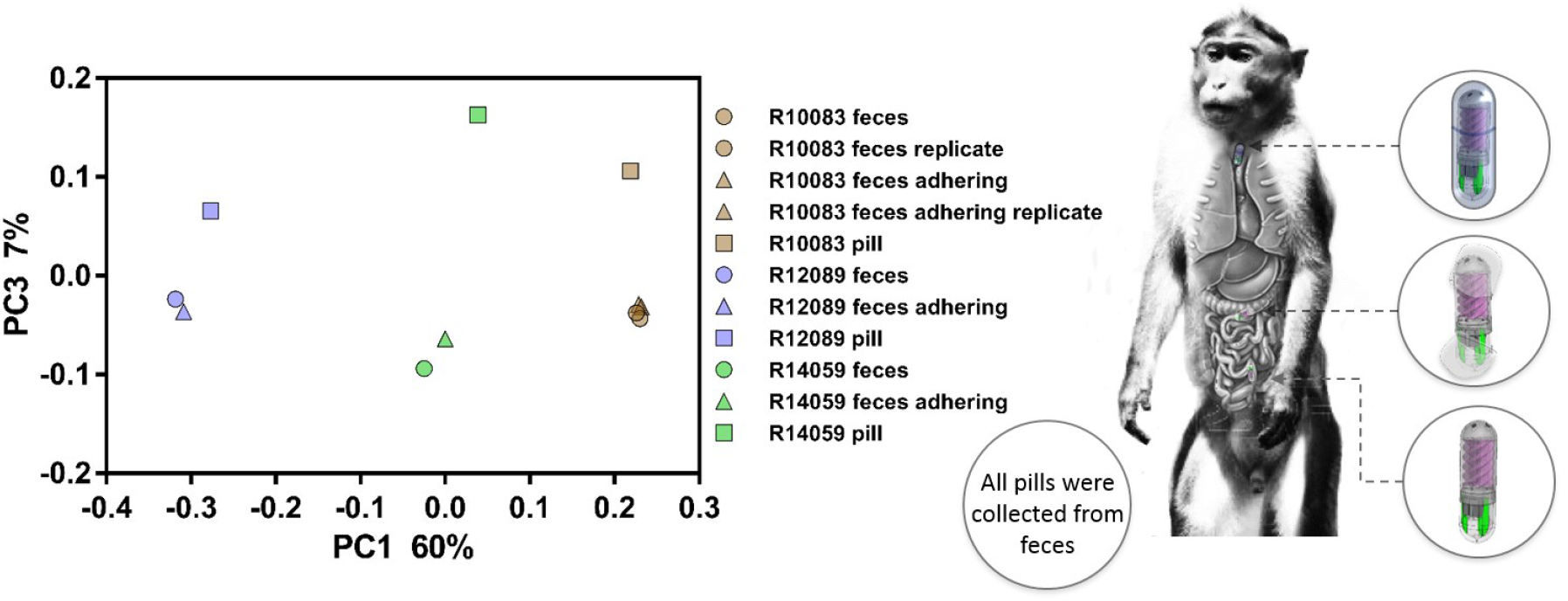
Principal Coordinate Analysis of bacterial microbiome sampled by pills orally administered to 3 macaques. Samples were extracted from the pills as described in Materials and Methods. In addition, DNA extracted from the feces and from feces adhering to the outside of the pills were sequenced to assess to what extent the material sampled by the pills differed from the feces. The colors represent the animal as indicated in the key with a 6-digit alphanumeric code. Symbol indicates origin of sample; square, pill’s collection channels, circle, feces; triangles, feces adhering to the outside of the pill. Identical symbols of the same color are technical replicates obtained by amplifying, barcoding and sequencing a sample twice. The distance between replicates is a measure of technical variation, i.e., variation introduced by sample processing and by sequencing.

We used Linear Discriminant Analysis to identify Operational Taxonomic Units (OTUs), which differ significantly in abundance between the pill’s collection channels and the feces.^[20]^ The 3 samples collected by the pills were highly enriched in Firmicutes belonging to the class Clostridia, order Clostridiales (Supplementary Table 2). Out of 32 OTUs significantly enriched in the pill samples, 21 (65.6%) belonged to this order, whereas in the fecal samples collected from the ground or adhering to the outside of the pills only 1/9 OTUs (11.1%) belonged to this taxon. The association between Clostridia classification and samples origin is statistically significant (Fisher exact test, p=0.003). Because the small intestine is difficult to access, little information is available on microbial populations residing in more proximal sections of the GI tract. However, the genus *Clostridium* is known to be more abundant in the small intestine than in the colon.^[21,22]^ The high abundance of Clostridiales OTUs in samples collected by the pills is consistent with active sampling as the pills travels through the macaques’ GI tract. The complete taxonomic profile of the samples recovered from the pills, from the feces and from fecal material adhering to the outside of the pills is shown in Supplementary Table 3.

### Towards spatially selective sampling

The *in vivo* and *in vitro* studies validated the pills functionality in sampling the gut environment with different anatomy and conditions. Our results show that the bacterial populations recovered from the pills’ microfluidic channels closely resemble the bacterial population demographics of the microenvironment to which the pill was exposed. Most importantly, it showed the ability of the pill to sample the regions of the gut upstream of the colon with quite distinct microbiome population compared to the feces. For a more spatially targeted sampling, one can immobilize the pills using an external magnet where more of the sample can be collected from a targeted region. We performed *in vitro* studies to validate the magnetic hold functionality for spatially selective sampling using colored dye as a model of GI fluid. The results reported in supplementary successfully validate our hypothesis (see supplementary Figure 4 and Supplementary Table 4). *In vivo* animal studies with magnetic hold for spatially selective sampling will require large magnets, *in vivo* imaging and visualization and other infrastructure – this is the basis for future work.

## Materials and methods

### Ingestible osmotic pill fabrication

The pill sampling head and the bottom salt chamber are 3D printed with *Form 2* 3D printer (Formlab Inc., Somerville, MA, USA) using high temperature resin also provided by Formlab. The resin is translucent and tough after curing in the FormCure photo-curing device (Formlabs Inc., Somerville, MA, USA) for 60 minutes. A reverse osmotic semipermeable membrane (GE Osmonics flat sheet membrane, SterliTech Co., Kent, WA, USA) was used to construct the osmotic pump. The membrane comprises of 5 μm thick woven cellulose acetate fibers (see Figure 1e). A circle of 6mm in diameter was cut from the membrane sheet using a laser cutter. Then, the membrane was placed on the back of the pill head. High temperature resin was cured on the periphery of the membrane to firmly affix the membrane in that position. A square neodymium magnet of 3 mm×3 mm×1.5 mm was placed in a specially designed cavity inside the salt chamber but isolated from the surrounding by HTR. Green fluorescent dye (part number: 37943991 by DecoArt®, Stanford, KY, USA) was added into two tube-like reservoirs in the salt chamber, dried and sealed by drop casting with a small amount of high temperature resin added into the containers. The salt chamber was filled with calcium chloride powder before the salt chamber and the sampling head were assembled together. The fabricated osmotic pill is shown in Figure 1b.

### Sample extraction

The inlet port that allows priming and extracting the collected sample after recovery from the feces was marked with a black dot. This port is directly connected to one of the helical channels. The four helical channels are connected at the bottom to the collection chamber. Sample recovery is achieved by applying suction at this port, or by centrifugation of the pill in a head-down position. Either method extracts the sample out of the corresponding helical channel, the collection chamber and then the other 3 helical channels in that sequence. A 100 μL pipette tip that easily fits into this marked inlet is used for this task Extraction is achieved using a pipette or a syringe. The process is shown in supplementary video 1. The centrifugation procedure is described below (section *In Vitro Experiments)*

### Motile bacteria culture

Fluorescent wild-type *Bacillus subtilis* (strain SG67 GFP) bacteria were cultured by inoculating 5 mL of Cap Assay Minimal (CAM) motility medium with cells obtained from a frozen glycerol stock solution.^[13]^ Cells were grown overnight at 37°C while shaking at 250 rpm until the optical density reached *OD*_600_ = 0.1. These cells have 1 μm×3 μm elongated bodies and swim at a speed of *U* = 40 – 65 μm.s^-1^.^[13]^

### Imaging and analysis

Epiluorescence imaging was performed on an inverted microscope (Nikon Ti-E) with a 10 × objective to take snapshots of the cells in the samples with a Zyla sCMOS camera (Andor Technology). Image analysis was performed to enumerate the bacteria by intensity thresholding in ImageJ.^[23]^

### Particle and bacterial concentration measurement

To quantify the concentration of the bacteria/particles collected by the pill over time, these steps were followed: 100 μL of the sample extracted from the pill using a pipette were dispensed on a glass slide; two spacers of 100 μm height where placed on either edges of the glass slide; another glass slide were placed on the top of the sample to confine the liquid; the sample was then imaged and the number of bacteria/particles on the surface were counted using the ImageJ software in the field of view. Three such images were taken over the entire glass slide. The average number of bacteria/particles per field divided by the visible area for the view, and the particle concentration expressed as number per mm^2^.

### In vitro experiments with live bacteria and fecal slurries

Experiments in which pills were immersed in suspensions of live bacteria were conducted to assess the ability of the pill’s osmotic pump to draw bacteria into the collection channel. To assess the effect of time on the ability of the osmotic pump to aspirate from the environment, a live suspension of each of the following bacterial species was prepared; *E. coli* strain K12, *B. subtilis* strain 168 BFA and *L. rhamnosus* (BEI Resources, strain LMS2-1, cat # HM-106). *E. coli* and *B. subtilis* were grown overnight in LB broth (Fisher BioReagents, cat. # BP1426-500) and *L. rhamnosus* in MRS broth (Fluka Analytical, cat. # 69966), respectively. The concentration of Colony Forming Unit (CFU) of each culture was estimated by plating 10-fold serial dilutions on agar plates. Bacterial suspensions were prepared in phosphate buffered saline at a concentration of 10^8^ CFU/ml and kept in motion with a magnetic stirrer. Pills without enteric coating were immersed sequentially for 4 hr in each suspension as follows: pill 1, 4 hr each in *E. coli*, *B. subtilis* and *L. rhamnosus.* Pill 2 and pill 3, 8 hr in *E. coli* followed by 4 hr in *L. rhamnosus.* To recover the content of the collection channel, each pill was introduced with the channel opening facing down into a 1.5 mL microcentrifuge tube and spun at 11,000 x g for 5 min. The pill was removed and the entire volume of approximately 200 μL recovered from the pill processed for DNA extraction. The same sample recovery method was used in all *in vitro* and *in vivo* experiments described below.

A second *in vitro* experiment was conducted to confirm that uptake of sample requires the action of the osmotic pump. Standard pills with CaCl_2_ in the salt chamber as shown in Figure 1b and pills lacking salt were immersed in parallel in fecal slurry. No enteric coating was used. The pills were kept in motion on a magnetic stirrer in 200 mL glass beakers containing fecal slurry. The experiment was conducted at room temperature.

A third *in vitro* experiment was undertaken to further assess the function of the pill when it is immersed in fecal slurry. Specifically, we investigated the level of contamination of the sample recovered from the collection channels with material adhering to the outside of the pills. Fecal slurry was prepared as described above for the second experiment. Two pills were immersed in each of the two slurries (cat or mouse) and retrieved after 24 hr. Thereafter each pill was briefly immersed in the opposite slurry (mouse or cat), rinsed with distilled water and dried.

### Experiments in pigs and macaques

Weaned pigs approximately 3 weeks of age were be purchased from the Cummings Veterinary School campus and housed in pens according to IACUC guidelines (protocol number G2018-03 Tufts University). The pills’ osmotic pump was activated with distilled water and the collection channels primed with water. The pills were enclosed in an enteric capsule (Size 00 white empty enteric coated capsules from CapsulCN International CO., Ltd, China) immediately prior to oral administration. The pill was inserted into a pill “gun” to introduce the pill directly into the pig’s esophagus, preventing regurgitation (see Supplementary Figure 5). Two experiments were conducted in pigs. In experiment 1, two pills were recovered from the stomach of the pig post-euthanasia. In experiment 2 one pill was recovered from the feces (see Supplementary Figure 6).

Experiments in non-human primates (see supplementary Figure S7) were conducted at the Biomedical Primate Research Center (BPRC) in Rijswijk, The Netherlands. Eight healthy rhesus macaques (*Macaca mulatta*) were selected from a group of animals that received a physical examination for a non-related scientific project (permit AVD5020020186346). This specific part was approved by the Animal Welfare Body (IvD 014A) and consisted of 6 males and 2 females, aged between 4 and 8 years. All animals were born and raised in the BPRC colony. Upon selection for studies, the animals were pair-housed in the experimental facility in primate cages of 1×2×2 m provided with bedding and enrichment. The animals were fed with commercial monkey pellets supplemented with vegetables and fruit, drinking water was provided *ad libitum.* For the physical examination, animals were sedated with 10 mg/kg Ketamine hydrochloride. The pill was manually placed into the pharynx of the animal and guided from the upper part of the esophagus to the stomach with help of a feeding tube. The animals were examined for 48 h and 4 times per day for stool passing by, observation and all fecal material was collected and palpated. Four pills were recovered from the fecal material of 4 animals (2 males, 4 and 8 years old, 2 females 6 and 8 years old) within the first 24 h. The remaining animals were followed for another 48 h and fecal material was collected and checked, but no pills were found. The animals from which no pills were recovered were euthanized after 2 months as part of the scientific project for which they were assigned. During the necropsy, no pills were found and the gastro-intestinal tract showed no significant findings. Also an additional CT scan of one of the animals just before necropsy did not show any evidence of the remains of a pill. It can be assumed that the unrecovered pills were passed by the animals, but were missed in the check of the fecal matter.

### Molecular biology methods

Following recovery of the content of the collection channels, DNA was extracted in a QiaCube instrument using the QIAamp PowerFecal DNA kit (QIAGEN, cat. 12830-50) according to the manufacturer’s protocol. DNA was eluted in 50μL elution buffer and stored at −20°C. A previously described PCR protocol was used to prepare 16S V1V2 amplicons libraries for 16S amplicon sequencing.^[24]^ The quality of the 16S amplicons was assessed by agarose gel electrophoresis. Amplicons were pooled in approximately equal molar ratio, size-selected with a Pippin Prep instrument (Sage Science, Beverly, Mass.) and sequenced in an Illumina MiSeq sequencer at the Tufts University Genomics core (tucf.org) using single-end, 300-nucleotide strategy. To control for technical variation introduced during library preparation and sequencing, each library included two replicates of two randomly chosen samples. Replication involved processing duplicate fecal samples, amplified and barcoded individually.

Bacterial DNA was quantified by real-time PCR using the same V1V2 PCR protocol as used to construct the 16S libraries. The PCR was performed in a Roche LightCycler instrument. Crossing point (*C_t_*) values, i. e., the number of temperature cycles required to reach a predetermined concentration of PCR product, were determined from the PCR amplification curves using the instrument’s software. To convert Ct to relative DNA concentration (*C*), *C_t_* values obtained from a 10-fold serially diluted DNA sample were regressed on the *log(C)*. The DNA concentration of this solution was determined using a Qbit 3.0 fluorometer (Invitrogen Life Technologies). The linear regression equation determined from these data was

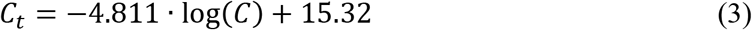

which was used for converting *C_t_* to DNA concentration (*C*).

### Bioinformatics analysis

Bioinformatics analysis of 16S sequences was performed in *mother* essentially as describe.^[24–26]^ Briefly, random subsamples of 5000 sequences per sample were processed using a sequence processing pipeline designed to generate pairwise weighted UniFrac distances between samples.^[27]^ Distance values were entered into a distance matrix and visualized by PCoA. PCoA plot were computed in GenAlEx.^[28]^ Sequences were taxonomically classified using the method described by Wang et al^[29]^ in *mothur* (same ref as [25]). The minimum bootstrap value for taxonomic assignment was set at 70%.

To convert relative 16S sequence abundance value for each bacterial species obtained in *In vitro* experiment 1 to relative abundance of bacteria, relative sequence abundance data were normalized against the number of rRNA operons present in the genome of each bacterial species. This calculation was necessary because the number of rRNA operons varies by species and sometimes also between strains of a same species. The number of operons obtained from the University of Michigan rrnDB database at https://rrndb.umms.med.umich.edu/ is 7 for *E. coli* K12, 10 for *B. subtilis* 168 BFA and 5 for *L. rhamnosus* LMS2-1.^[30]^

## Acknowledgments

Office of Naval Research (ONR) primarily sponsored this work under the grant N0014-16-1-2550 (Program manager: Dr. Linda Chrisey). B.C.M.O and G.W. were supported by the National Institute of Allergy and Infectious Diseases (grant # 5R21AI125891). A.D and J.G would like to acknowledge the support of National Science Foundation (CBET-1511340, CAREER-1554095). Our thanks to Boudewijn Ouwerling and Don Girouard for expert assistance with animal experiments and to Dr. Albert Tai and the Tufts Genomics Core Facility staff for high-throughput sequencing.

## Author contributions

S.S. invented the concept of the pill. H.R.N and S.S conceptualized the design of the pill and co-wrote the manuscript. H.R.N fabricated the pills and performed *in vitro* and some *ex vivo* studies. H.R.N ran the simulations and prepared figures for the manuscript. G.W. led the animal studies effort and co-wrote the manuscript. S.T. supported the animal studies in pigs. I.K and J.A.M.L performed *in vivo* animal studies in monkeys. B.C.M.O. and G.W. performed *in vivo* studies in pigs and *ex vivo* experiments and did data analysis and interpretation for all animal studies. H.R.N. and A.S. performed the magnetic hold test experiment. A.D., J.G. and H.R.N. performed *in vitro* sampling studies with motile bacteria. A.S. also helped prepare some figures. All authors discussed the results and commented on the manuscript.

## DNA sequence accessions

16S DNA sequence data were deposited in the European Nucleotide Archive under accession numbers PRJEB30052 (Pig experiment and *In vitro* experiment 3) and PRJEB32383 (Primate experiment and *in vitro* experiment 2)

## Supplementary Information

### 1. Estimation of flow rate of the passive osmotic pump

Two sets of experiments were carried out to estimate the flow rate over time through the membrane under different backpressure conditions. Osmotic flow depends on the differential pressure generated due to ionic concentration gradient across the membrane. In the pill, the pressure is higher on the top side, where the ion concentration is lower, and it is lower on the bottom side where the ion concentration is high (side facing the salt chamber). This pressure difference causes the water to flow across the membrane from the side with lower ion concentration towards the side with a higher ion concentration. To quantify the osmotic pressure, we measured the pressure at the discharge pore needed to stop the osmotic flow. Supplementary Figure 1a shows the pressure measurement setup. Here, the membrane separates two reservoirs (red and blue). The bottom reservoir is filled with deionized water containing red food dye and the top reservoir is filled with a saturated CaCl_2_ solution containing blue food dye. If the water level in both the reservoirs is identical, no external pressure would be applied on the membrane. However, by increasing the level of water in the blue reservoir, an external pressure difference is applied to the membrane. This pressure difference can be calculated using the relation Δ*P* = *ρg*Δ*h* (*ρ* is the density of saturated salt water, *g* is the gravity and Δ*h* is the height difference between the reservoirs). The flow rate was calculated based on the observed increment in the level of the top reservoir over 1 hr, multiplied by the cross-sectional area. We observed that the osmotic flux through the membrane linearly decreases as we increase the external pressure (Supplementary Figure 1a). After applying a 1.5 kPa pressure difference, a decrement in the osmotic flux of about 25% was observed. Based on its linear trend, we interpolated the pressure causing the flux of water to stop at ~4 kPa. The flow rate through the membrane assembled on the pill was also measured over time (Supplementary Figure 1b). Initially, the flow rate was ~2.5 μL/hr and remained nearly constant for almost 48 hr. The flow rate fell to around ~0.5 μL/hr after four days.

**Supplementary Figure 1.**
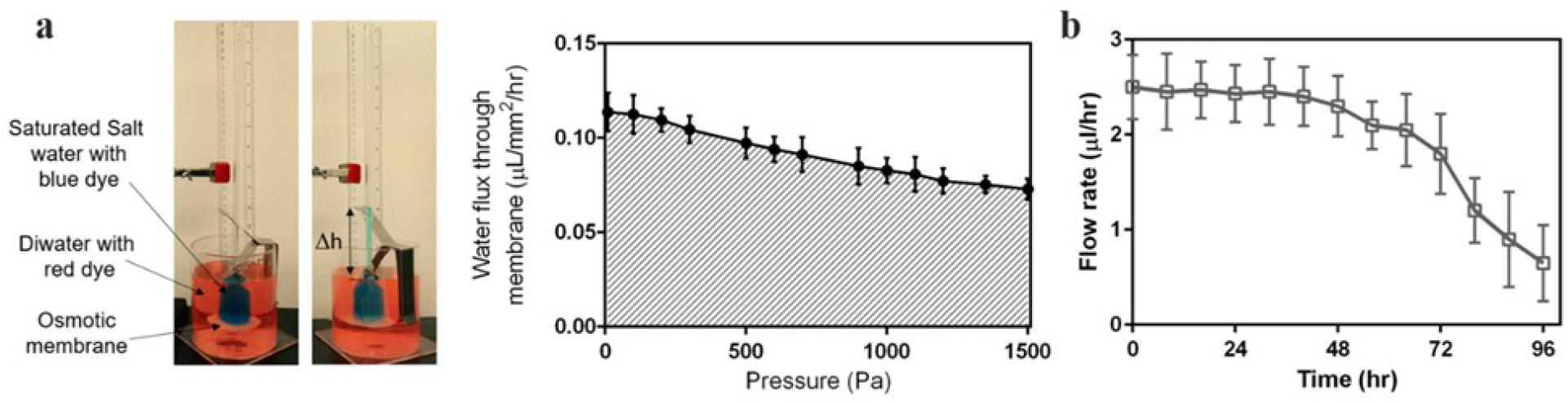
Fluid flow characterization of the osmotic membrane and the pill. a) osmotic characterization test setup and water flux under different back pressure. b) osmotic membrane performance through time.

### 2. Numerical simulation of osmotic flow through discharge hole

The discharge hole in the salt chamber at the posterior end of the pill is important because it facilitates the discharge of any excess water in the salt chamber. A series of computational fluid dynamics simulation (COMSOL Multiphysics®, Burlington, MA, USA) was carried out to study flow through the discharge hole. Two different hole sizes (100 μm and 50 μm diameter) were examined and the maximum velocity of the fluid estimated. See Supplementary Figure 2 for simulation results. Numerical results show that the maximum fluid velocity for the 50-μm hole is 0.6 mm/s, which is 4.28 times higher than for hole of 100 μm size. A higher continuous flow rate at the discharge point has two advantages: first, it reduces the diffusion from the environment into the salt chamber, essentially acting as a one-way valve; second, a high flow rate reduces the chance of blockage at the discharge hole by solid debris present in the gut.

**Supplementary Figure 2.**
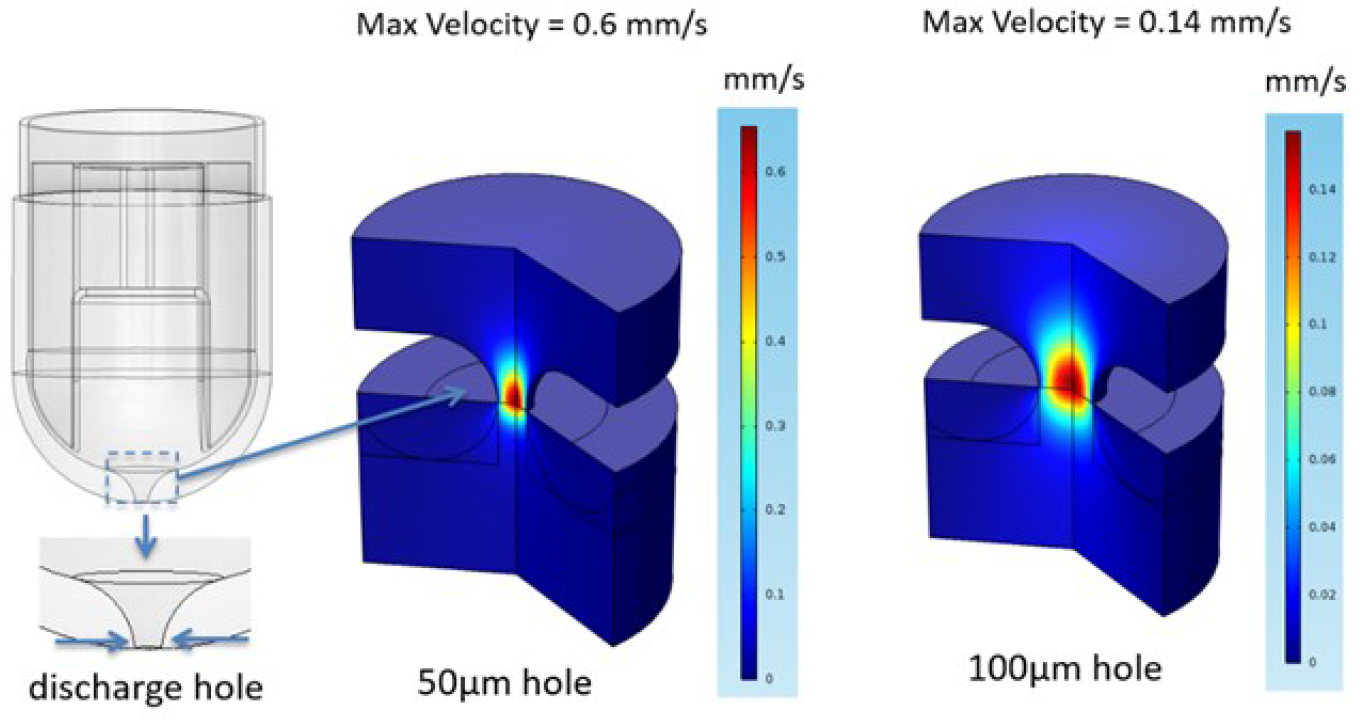
Numerical simulation of flow through different size of discharge hole.

### 3. Pill mobility in the gut (ex-vivo study)

A 1 m portion of small intestine was used to carry out the test. The setup used for the ex-vivo test is shown in Supplementary Figure 3. The intestine was emptied of its content and placed horizontally in 1 L of water. The pill was inserted into one of the intestine’s opening. The opening was then covered with silicone tubing with outer diameter of 6 mm using a clipper to create a tight connection. The other end of the intestine was left open. An external peristaltic pump was used to simulate a peristaltic flow inside the GI tract. To mimic highly viscous gut content, a solution of 5% cellulose hydrogel (w/v), rhodamine (for visualization) and water was used. The fluid had a dynamic viscosity of approximately 100 mPa.s. Finally, three different flow rates for the pumping solution were tested, and the movement of the pill under these flow conditions was measured. The experiment terminated when the rhodamine-loaded hydrogel emerged at the other end of the intestine. We repeated this test three times with the same tissue.

**Supplementary Figure 3.**
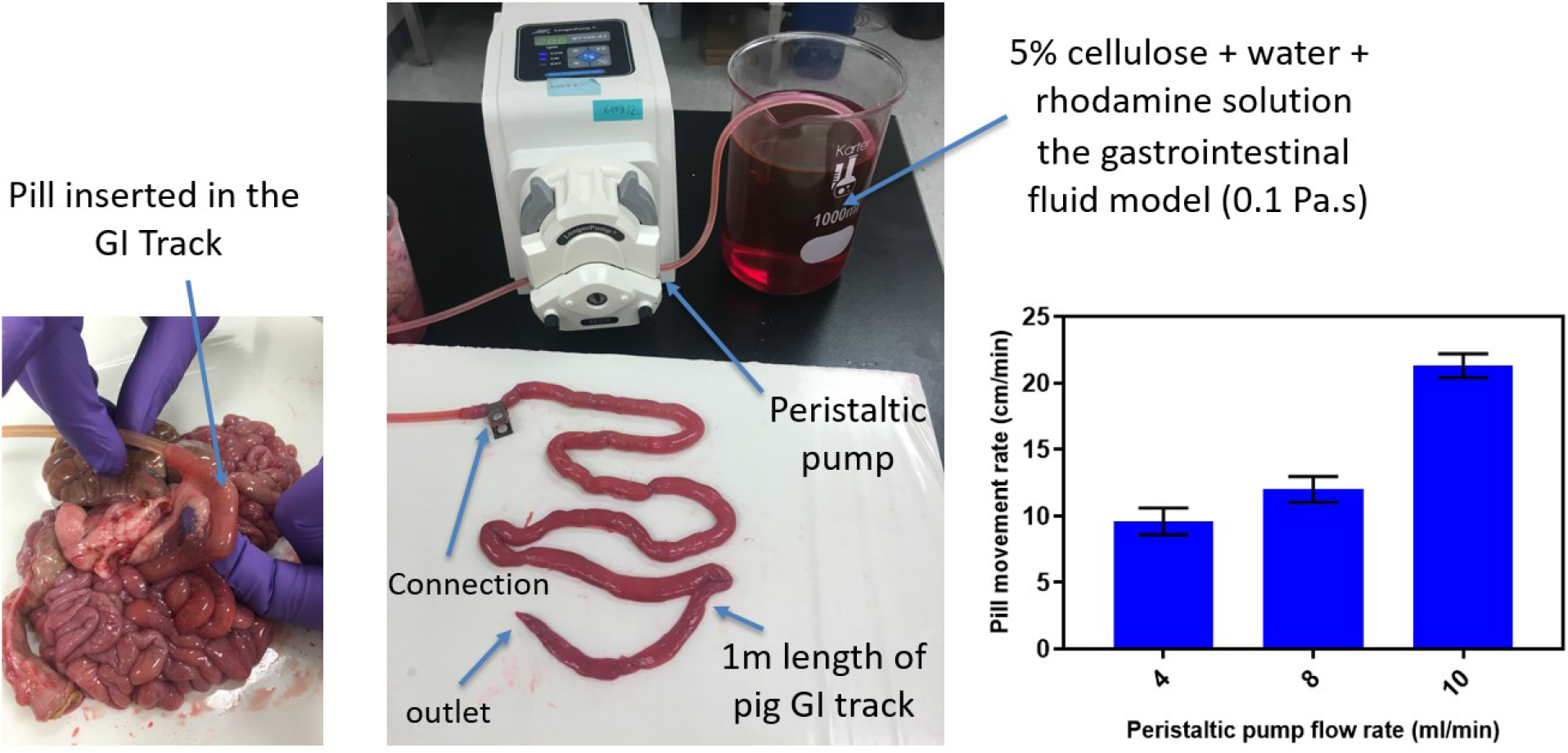
Ex-vivo test of pill mobility inside the pig intestine. A simulated gut media was pushed through the intestine with an external peristaltic pump to mimic natural peristalsis.

### 4. Diffusion coefficient of micro-particles

To validate that the diffusion effects can be neglected, we calculated the diffusion coefficient for micro-particles in the stated conditions using the Einstein-Smoluchowski relation,

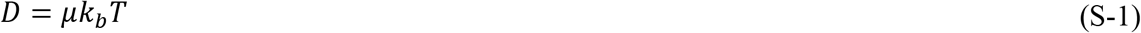

where *μ* is the mobility coefficient expressed as the ratio of the particle’s terminal drift velocity to an applied force, *μ* = *v_d_*/*F*), *k_b_* is Boltzmann’s constant and *T* is the temperature in Kelvin scale.

At low Reynolds number, the Stokes-Einstein relation for spherical particles of diameter *d* is,

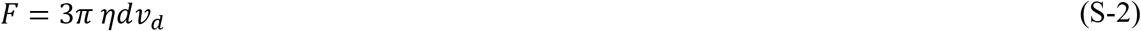

where is the dynamic viscosity of the medium. Thus the Einstein-Smoluchowski relation simplifies to the Stokes-Einstein relation,

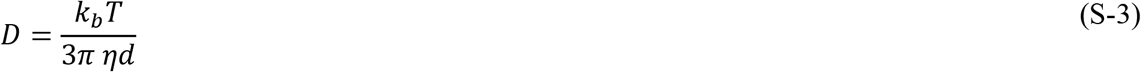

Based on the Stokes-Einstein relation, the diffusion coefficient of a spherical particle of 10 μm in diameter at the test temperature of 37°C in a fluid with the viscosity of 160 mPa.s is 2.8 × 10^-4^ μm^2^/s, which is very small and can be neglected in our study.

### 5. Taxonomic classification of samples from pigs and primates

Supplementary Table 1.

Taxonomic classification of 16S sequences from 7 samples extracted from two pills recovered from pig feces and from 3 GI tract organs. Sequences were classified using program *classify.seqs* in mothur [25]. The headers are color-coded to indicate the origin of the sample as indicated in Figure 4.

Supplementary Table 2.

Bacterial OTUs significantly differing in relative abundance in samples originating from the pills’ collection helical channels (square symbols in Figure 5) as compared to samples collected directly from the macaques’ feces or adhering to the outside of the pills (circle and triangle symbols, respectively, in Figure 5). OTUs enriched in sample collected by the pills are indicated with red font. OTUs enriched in the feces are indicated with green font.

Supplementary Table 3.

Taxonomic classification of 11 primate experiment samples. Column headers are color-coded as in Figure 5. Samples “Feces” and “Adhering Feces” in columns E, F and G, H, respectively, are replicates to control for technical variation.

### 6. Magnetic hold test

The pill has a micro-magnet that serves two purposes: first, to hold the pill in place at a desired section of the GI tract to acquire more sample from a specific location; second, to facilitate recovery of the pill following excretion in the feces. An *in vitro* test was designed to validate the local sampling capability of the pill in three different environments. Instead of creating three different chambers each with a different environment, we created an equivalent setup where there is just one chamber and we pump in three different test solutions mimicking three different *in vivo* environments (see Supplementary Figure 4). The setup is shown in Supplementary Figure 4a. Initially, the absorption spectrum of each individual dye and a mixture of the three dyes, was captured to ensure that the interaction between dyes is negligible (see Supplementary Figure 4b). In the first test, we filled the feeding reservoir with yellow dye test solution and applied a flow rate of 2 mL/sec (same for all tests) to move the solution in the testing chamber for 90 min. The chamber was then filled with red dye for 30 min and then with blue dye for another 30 min. After finishing all three steps, the pill was recovered and 100μL sample extracted from the pill’s collection channels using a pipette. The sample was then mixed with 2 mL of distilled water and analyzed with a UV-Visible spectrophotometer. The resulting spectrum in Supplementary Figure 4c shows that the absorption of the yellow dye is higher relative to the red and blue dyes. We also decomposed the spectrum response to find the individual components from individual dyes in the sample (Supplementary Figure 4c). In test 2, we put the pill in yellow dye test solution for 30 minutes, red dye for 90 minutes and blue dye for 30 minutes. As we could expect from the previous experiment, the red dye has higher peak than the yellow and blue dyes as seen in Supplementary Figure 4d. In test 3, we put the pill in yellow dye test solution for 30 minutes, red dye for 30 minutes and blue dye for 90 minutes. The blue dye shows a higher peak than the other two dyes as shown in Supplementary Figure 4e. The tests are summarized in Table S1.

The purpose of the test was to validate the hypothesis that the pill can be held in a desired place using an external magnet, even in moving fluid, and that the pill will collect more sample from a region it is immersed in for more time. The experiment was performed in a plastic tube with an internal diameter of 2 cm and a thickness of 1.5 mm (see Supplementary Figure 4). An external neodymium magnet was used to hold the pill in place. Peristaltic flow was induced in this chamber using an external peristaltic pump. To mimic sampling in three different areas of the GI tract, we pumped three different aqueous test solutions. The time the pill is exposed to each of the test solution dictates how much of that solution is aspirated into the helical channels. Dyes were used to visually identify the three different test solutions in which the pills were suspended. In this setup, we pumped two of three test solutions for 30 min which mimics the transit regions of the GI tract, and the remaining third for 90 min which mimics the “hold time” in the desired location of the GI tract (see Table S1). The absorption spectrum of the sampled contents recovered from the pill following the experiment was analyzed using a UV-Vis spectrum analyzer.

The setup for this *in vitro* test is shown in Supplementary Figure 4a. Each feeding reservoir contains one of the three test solutions. A tube was connected from the peristaltic pump to the testing chamber in which the pill was placed. Another tube connects the outlet of the testing chamber to a collection reservoir. We used a grade N52 Neodymium magnet (TotalElement®, Centennial, CO, USA) 25.4 mm×25.4 mm×25.4 mm to hold the pill in the middle of the testing chamber. The magnet was placed 10 cm away from the testing chamber. We chose yellow (Brilliant Yellow from Sigma-Aldrich, Natick, MA, USA), red (metallic red from Chefmaster®, Fullerton, CA, USA) and blue (metallic blue from Chefmaster®, Fullerton, CA, USA) dues to distinguish the three test solutions. These dyes have different spectral responses and more importantly, once mixed, their spectrum signature is not influenced by the presence of other dyes. We mixed 100 μL of each dye separately with 2 mL of distilled water and analyzed the solutions with an Evolution 220 UV-Visible Spectrophotometer (from Thermo Fisher Scientific, Waltham, MA, USA) to obtain the spectral signatures. The scanning range was between 340 nm and 700 nm, showing peaks at 403 nm, 499 nm and 629 nm for the yellow, red and blue dye, respectively. Note that the dyes have a broad spectrum around their peak absorption wavelength. We also mixed 100 μL of each dye with 2mL distilled water and extracted the spectrum of the mixed solution as seen in Supplementary Figure 4b. This analysis validated that the dyes have negligible interaction and that the superposition assumption is valid.

**Supplementary Table 4.** magnetic hold test in dye solutions

|  | Yellow dye (1<sup>st</sup>) | Red dye (2<sup>nd</sup>) | Blue dye (3<sup>rd</sup>) |
| --- | --- | --- | --- |
| **Test 1** | 90 min | 30 min | 30 min |
| **Test 2** | 30 min | 90 min | 30 min |
| **Test 3** | 30 min | 30 min | 90 min |

**Supplementary Figure 4.**
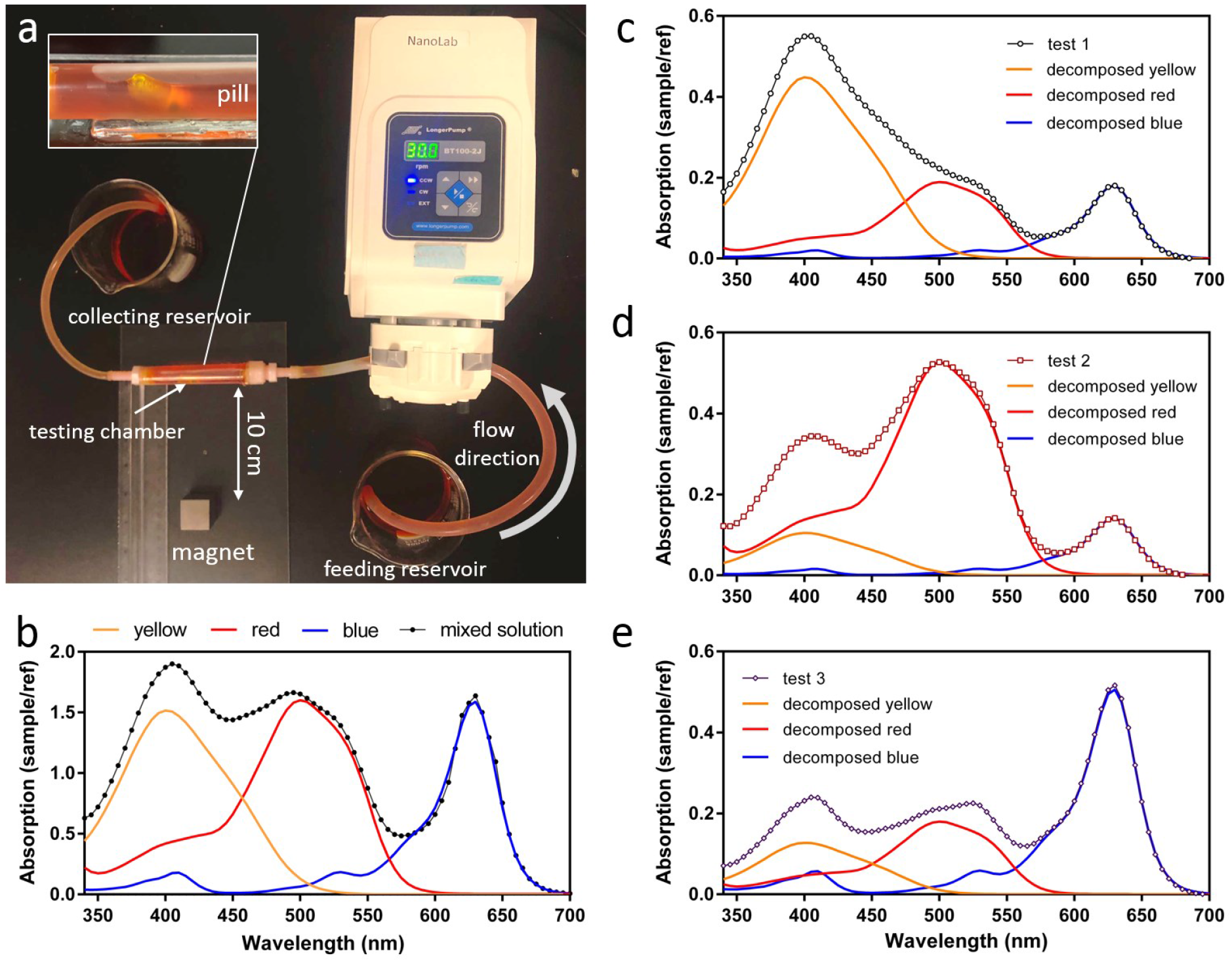
Validation of “trap and hold” sampling using a magnet. a) Experimental setup, b) spectrum of three different dye solutions and their mixture, c) test1 d) test2, a) test3 results. Results indicate that the content of the pill strongly correlates with the solution it is exposed to.

### 7. In vivo test – oral administration

**Supplementary Figure 5.**
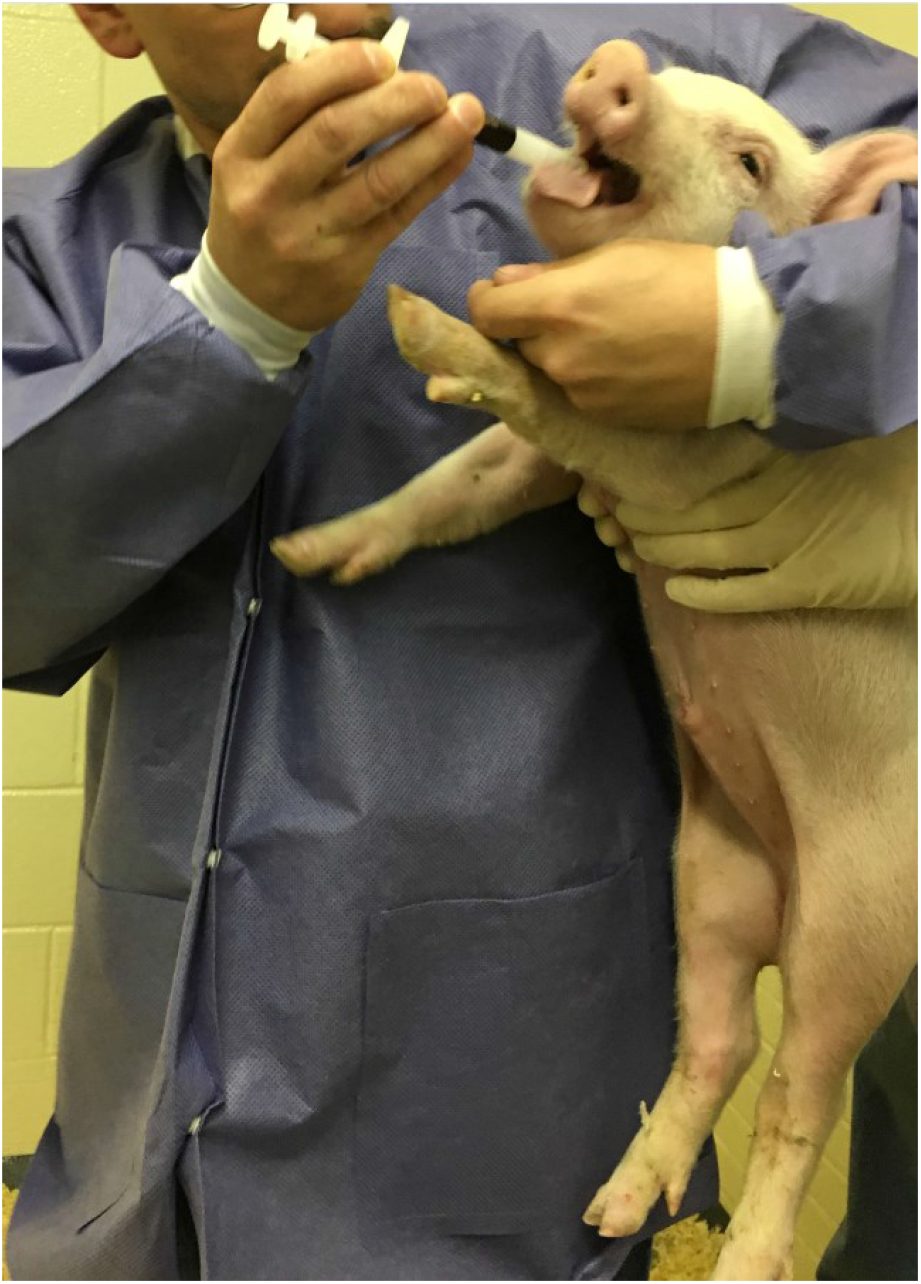
Oral administration of the pill for In-vivo test. The pill was inserted into a pill “gun” to introduce the pill directly into the pig’s esophagus, preventing regurgitation

**Supplementary Figure 6.**
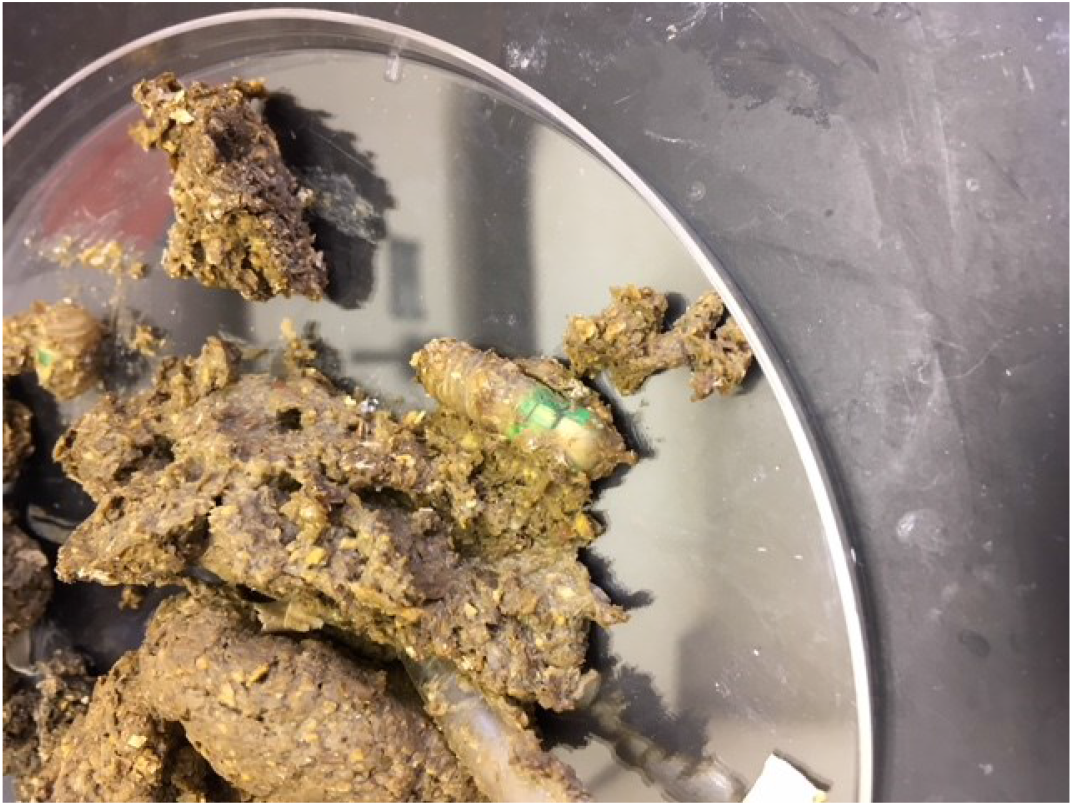
Extracted pill from pig feces in the in vivo experiment.

**Supplementary Figure 7.**
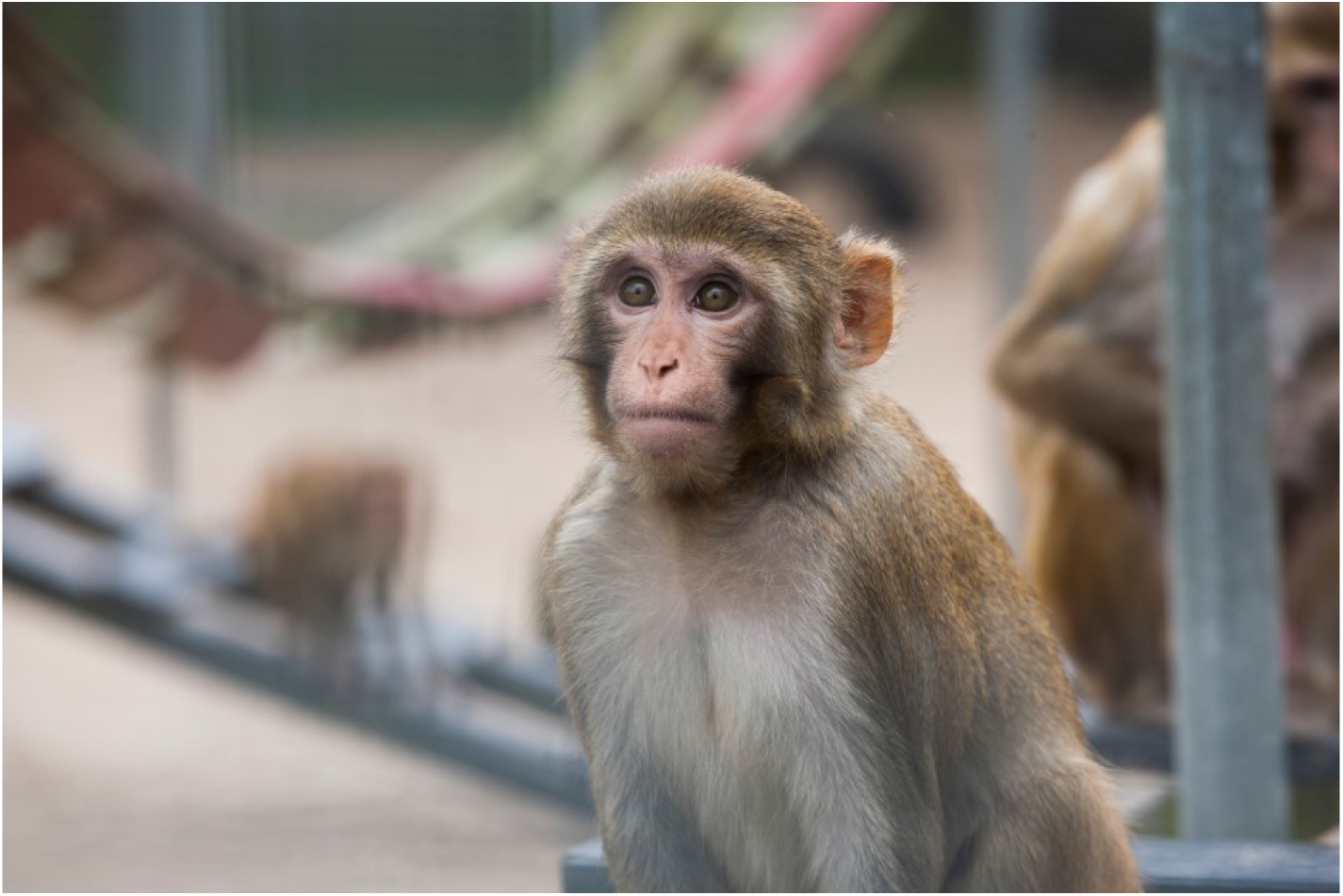
Rhesus monkey.

